# Tolerance-conferring defensive symbionts and the evolution of parasite virulence

**DOI:** 10.1101/2022.12.05.519091

**Authors:** Cameron A. Smith, Ben Ashby

## Abstract

Defensive symbionts in the host microbiome can confer protection from infection or reduce the harms of being infected by a parasite. Defensive symbionts are therefore promising agents of biocontrol that could be used to control or ameliorate the impact of infectious diseases. Previous theory has shown how symbionts can evolve along the parasitism-mutualism continuum to confer greater or lesser protection to their hosts, and in turn how hosts may coevolve with their symbionts to potentially form a mutualistic relationship. However, the consequences of introducing a defensive symbiont for parasite evolution and how the symbiont may coevolve with the parasite have received relatively little theoretical attention. Here, we investigate the ecological and evolutionary implications of introducing a tolerance-conferring defensive symbiont into an established host-parasite system. We show that while the defensive symbiont may initially have a positive impact on the host population, parasite and symbiont evolution tend to have a net negative effect on the host population in the long-term. This is because the introduction of the defensive symbiont always selects for an increase in parasite virulence and may cause diversification into high- and low-virulence strains. Even if the symbiont experiences selection for greater host protection, this simply increases selection for virulence in the parasite, resulting in a net negative effect on the host population. Our results therefore suggest that tolerance-conferring defensive symbionts may be poor biocontrol agents for population-level infectious disease control.

**Lay Summary:** Defensive symbionts – microbes that confer protection to a host against a harmful parasite – are found throughout the natural world and represent promising candidates for biological control to combat infectious diseases. Symbionts can protect their hosts through a variety of mechanisms that may prevent infection (resistance) or increase survival following infection (tolerance), yet our understanding of the ecological and evolutionary impact of defensive symbionts on parasites is limited. Moreover, few theoretical predictions exist for how defensive symbionts are likely to evolve in the presence of parasites, and for the net effect on the host population. Using a mathematical model where defensive symbionts reduce parasite virulence (harm to the host), we investigate the impact of their introduction on the evolution of parasite virulence, how selection increases or decreases host protection, and whether such symbionts are beneficial for the host population. We find that this form of defensive symbiosis always selects for higher parasite virulence and that it can cause the parasite to diversify into high and low virulence strains which specialise on different host subpopulations. Crucially, we show that the introduction of a defensive symbiont will always lead to a long-term reduction in host population size even if they are beneficial in the short-term. Together, our results show that defensive symbionts can have a strong impact on the evolution of virulence and that this form of host protection is not robust, indicating that tolerance-conferring symbionts are likely to be poor candidates for biological control of infectious diseases at the population level.

## Introduction

Defensive symbiosis, where an organism confers protection to its host from a natural enemy such as a parasite or predator is widespread in nature (reviewed in (Ford and King 2016)). For example, ants have long been known to defend acacia trees from herbivores (Belt, Thomas 1874) and various bacteria have been shown to confer protection directly or indirectly against bacterial and fungal parasites across diverse host taxa, including insects (Oliver et al. 2003; Cariveau et al. 2014), plants (Herre et al. 2007; Arnold et al. 2003), invertebrates (Gil-Turnes, Hay, and Fenical 1989; Gil-Turnes and Fenical 1992), and vertebrates (Lauer and Hernandez 2015; Heikkilä and Saris 2003). Protection can be conferred to hosts as resistance (preventing infection) or tolerance (increasing survival following infection) through a variety of mechanisms (Troha and Ayres 2022), including through interactions with the host’s immune system (Ford, Drew, and King 2022), interference competition through chemical defences - for example, *Streptococcus pneumoniae* can produce hydrogen peroxide to displace *Staphylococcus aureus* in the nasopharynx (Selva et al. 2009) – and resource competition or priority effects (Hancock, Sinkins, and Godfray 2011; Moreira et al. 2009). Defensive symbionts therefore have potential as agents of biocontrol, especially in the context of infectious diseases for therapeutic use (Bakken et al. 2011) or for population-level control (Utarini et al. 2021). The use of defensive symbionts should be approached with caution, however, as the nature and extent of protection conferred to their hosts is evolvable and they could alter both the ecological and evolutionary dynamics of hosts and parasites, potentially leading to unintended consequences.

Crucially, the protective relationship between a defensive symbiont and its host is not fixed; it may be context dependent, due to changes in the biotic or abiotic environment (González et al. 2021; King et al. 2016; Ashby and King 2017; Rafaluk-Mohr et al. 2018; Rogalski et al. 2021; Chamberlain, Bronstein, and Rudgers 2014; Lin and Koskella 2015), and it is subject to selection (King et al. 2016; Rafaluk-Mohr et al. 2022). For example, the removal of large herbivores can lead to the loss of acacia tree protection by ants (Palmer et al. 2008) and protective microbes such as *Enterococcus faecalis* reduce nematode fitness in the absence of *Staphylococcus aureus* but can be experimentally evolved to rapidly increase protection of their hosts when *S. aureus* is present (King et al. 2016). An organism may therefore be parasitic to its host in isolation, but may be protective – and may evolve to be more or less protective – when another parasite is present (Ashby and King 2017; Rafaluk-Mohr et al. 2018). Understanding evolution along the parasitism-mutualism continuum is therefore a key challenge for evolutionary biologists, especially in the context of the gut microbiome and infectious diseases. In particular, understanding the evolutionary robustness of host protection is particularly important when defensive symbionts are used as biocontrol agents, as their effectiveness will depend on both on the initial impact on the parasite, as well as the subsequent coevolutionary dynamics between host protection and parasite virulence.

The evolution of parasite virulence has long been a focus of theoretical studies of host-parasite systems (S. Levin and Pimentel 1981; R. May and Anderson 1983; Lenski and May 1994). Theoretical studies of host-associated communities have primarily focused on the effects of within- and between-host competition on the evolution of virulence (Alizon 2013; Frank 1992; 1996; Brown, Hochberg, and Grenfell 2002; R. May and Nowak 1995). By comparison, few theoretical studies have explored microbial evolution in the context of defensive symbiosis. Ashby and King (2017) explore how host protection evolves in the presence of a non-evolving parasite population, showing that conferred tolerance and resistance could readily evolve under a wide range of conditions, potentially leading to symbiont diversification into a highly protective strain and one that conferred no protection. This model was extended by Rafaluk-Mohr et al. (2018) to explore symbiont coevolution with the host, showing that the host becomes more mutualistic towards the symbiont at intermediate levels of protection. Nelson and May (2017) investigate the evolution of symbionts along the full mutualism-parasitism continuum when there is a shared cost of virulence. They show that the community of symbionts maintain mutualisms and evolve lower virulence when the shared costs are sufficiently low, but higher virulence may evolve when shared costs are high. Nelson and May (2020) extend this model to show that if increased defence is evolved by one symbiont, it may facilitate the reduction of virulence in both symbionts present, and in some cases cause pathogens to evolve towards mutualism. Together, these studies highlight the complex context-dependent nature of coevolution between mutualistic and parasitic symbionts. A key question, yet to be addressed by previous studies, is how do defensive symbionts drive the evolution of virulence in obligate parasites? Furthermore, how does the evolution of virulence affect the evolution of host protection, and what are the consequences for the host population?

Here, we use a mathematical model to address these questions. We explore the (co)evolution of parasite virulence and host protection – specifically, tolerance – by a defensive symbiont. Although tolerance can take many form (Rafaluk-Mohr et al. 2022), here we focus on mortality tolerance, whereby infected hosts that possess the defensive symbiont experience a lower mortality rate than infected hosts that do not possess the defensive symbiont. We first show how the introduction of a defensive symbiont always selects for greater parasite virulence, and that the defensive symbiont can induce the parasite to diversify into high and low virulent strains. We then show how the shape of life-history trade-offs associated with host protection affect the outcome of symbiont-parasite coevolution, and that this always results in a reduction in the host population size in the long term.

## Materials and methods

### Model

We consider a well-mixed population of hosts with two co-circulating microbes: an obligate parasite that increases host mortality and a defensive symbiont that may confer tolerance to infected hosts by reducing their disease-associated mortality rate. Hosts may exist in one of four states, where they harbour: no microbes (*H*), defensive symbionts only (*D*), parasitic microbes only (*P*) or both (*B*). New hosts are born at rate *ν* (*N*)(*a*−*qN*), where *N* = *H* + *D* + *P* + *B* is the total number of hosts, *a* is the maximum per capita rate of reproduction and *q* controls the strength of density dependent competition among hosts. All hosts, regardless of infection status, have a natural mortality rate *b*.

We assume that transmission is density-dependent, occurring at a baseline rate of 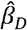 for the defensive microbe with a clearance rate of *γ*_*D*_, and *β*_*p*_ for the parasite with a clearance rate of *γ*_*p*_. There is no vertical transmission (all individuals begin life without either microbe), co-transmission does not occur (i.e., hosts must transition through one of the single-microbe classes to reach class *B*), and there is no long-lasting immunity. Both defensive and parasitic microbes increase the baseline mortality rate of the host, by *α*_*D*_ and 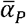, respectively. We assume that the parasite experiences a power-law trade-off between transmission and virulence (the additional microbe-induced mortality) such that 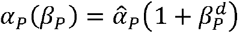 with *d* > 1 so that there are diminishing returns for increased virulence. Note that due to this positive correlation between parasite transmission and virulence, we will interchangeably refer to transmission and virulence evolution throughout. Defensive microbes may confer protection to parasitised hosts in the form of tolerance, *y* ∈ [0,1], such that the additional mortality rate for hosts with both microbes, *α*_*B*_ (*y*, β_*p*_), satisfies *α*_*B*_ (*y*, β_*p*_), ≤ *α*_*P*_ (β_*p*_),+ *α*_*D*_ (i.e. it is less than or equal to the sum of the additional mortality rates). However, the defensive microbe incurs a fitness cost when it diverts resources to protect a host, resulting in a reduction in its transmissibility such that 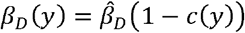, where *c*(*y*) is an increasing, non-linear cost function:

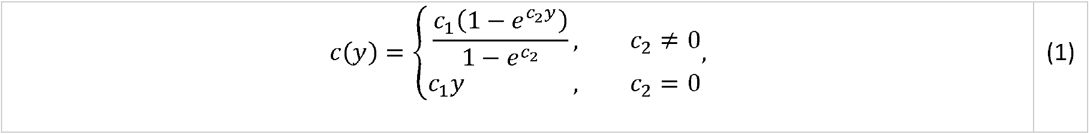

where ***c***_1_ ∈ [0,1] is the strength of the cost function, denoting the maximum reduction in transmission when tolerance is maximised at *y* = 1, and *c*_2_ controls the shape of the trade-off: when *c*_2_ > 0, conferring protection is increasingly costly (an accelerating trade-off), and when *c*_2_ < 0 conferring protection is decreasingly costly (a decelerating trade-off). Biologically, an accelerating (respectively, decelerating) trade-off corresponds to diminishing (increasing) returns, such that for a given increase in the cost, the amount of additional host protection decreases (increases) as it gets stronger. Thus, for accelerating trade-offs there is a relatively small initial cost but the costs become ever more extreme at higher levels of protection, whereas for decelerating trade-offs there is a relatively large initial cost but the additional costs become less extreme at higher levels of protection.

The ecological dynamics of monomorphic populations are shown schematically in Fig. 1 and are governed by the following ordinary differential equations (ODEs)

**Fig. 1:**
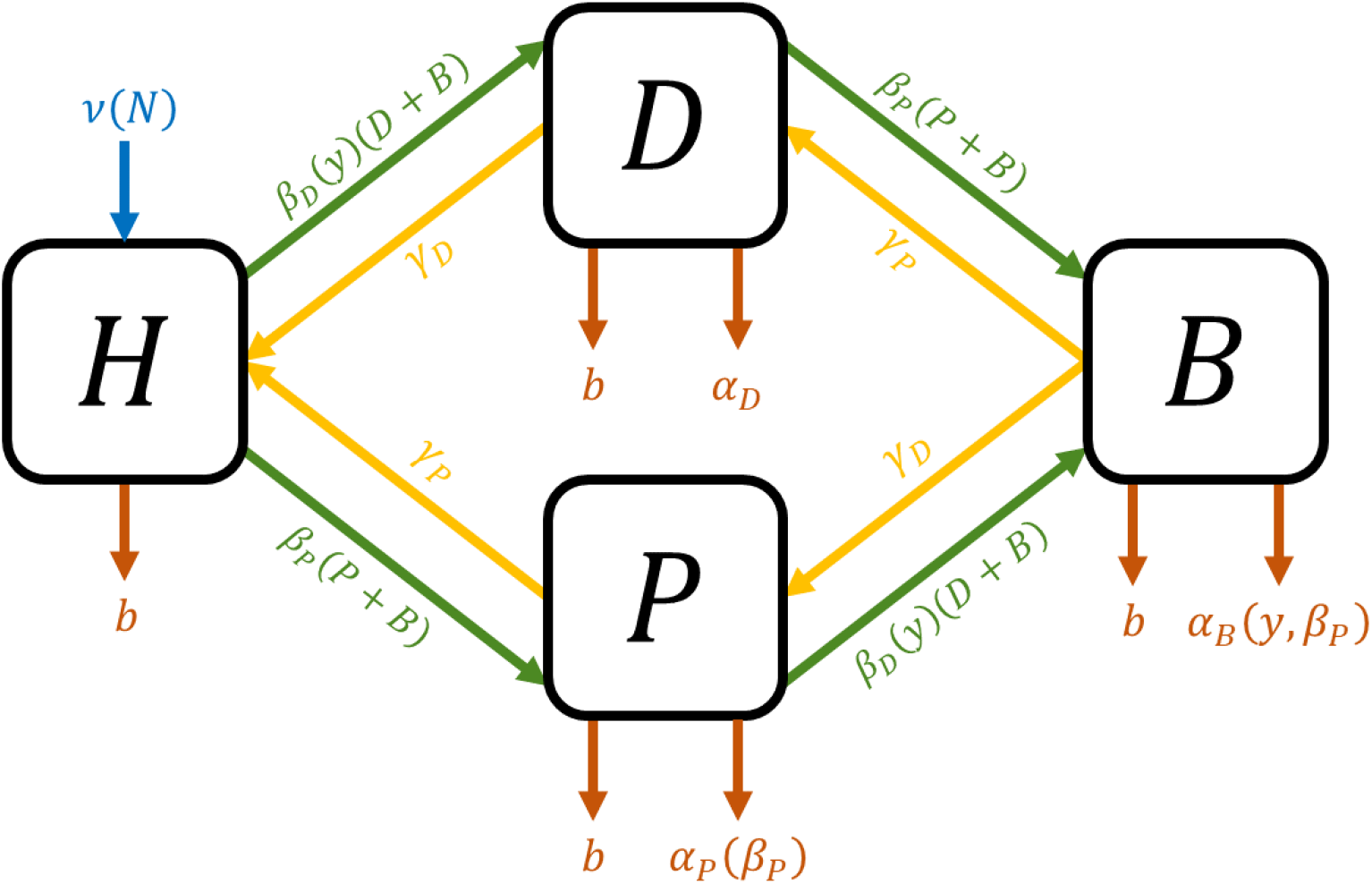
Model schematic. Arrows denote transitions into or out of states at the indicated rates: transmission (green), mortality (red), recovery/clearance (yellow) and birth of new hosts (blue).

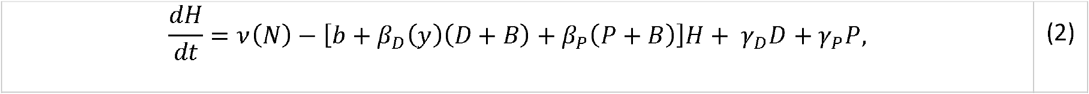

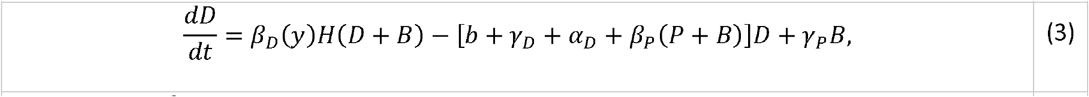

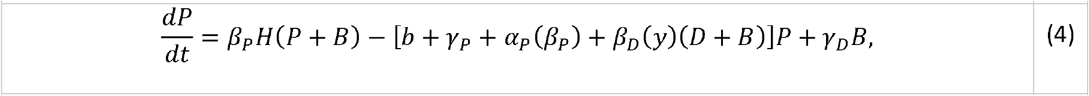

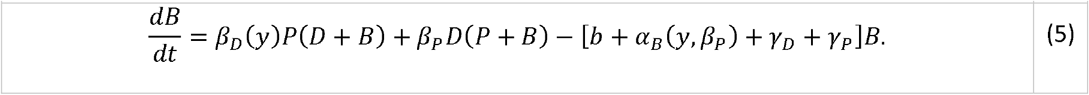

### Analysis

We employ evolutionary invasion analysis using a combination of numerical analysis and simulations to establish how parasite virulence (*α* _*p*_) evolves following the introduction of the defensive symbiont, and in turn how the defensive symbiont co-evolves to be more or less protective (*y*) following its introduction.

We use the next generation method (Diekmann, Heesterbeek, and Roberts 2010) (see *Supplementary Material*) to derive the invasion fitness for a rare defensive symbiont with protection *y*^*m*^ (denoted *w*_*D*_), or a rare parasite with transmission rate 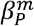 and virulence 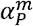 (denoted *w*_*p*_), when introduced into a population at equilibrium with resident traits 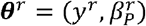:

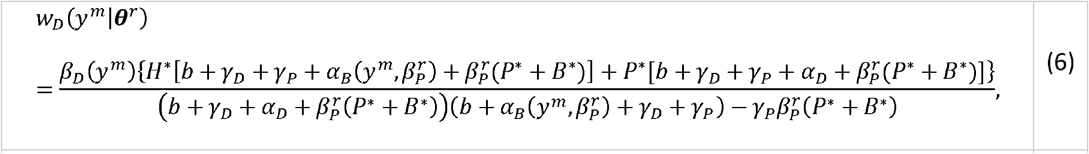

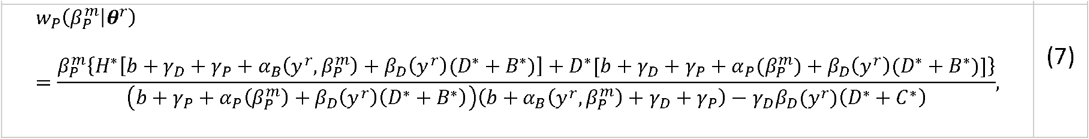

where each of the steady states (indicated with asterisks) are functions of the resident traits, for example *H*^*^ ≡ *H*^*^ (***θ***^*r*^). We are unable to obtain an analytical expression for these steady states, so we approximate them in our numerical analysis by simulating the ODE system for a sufficiently long period of time so that the population approaches its unique, locally asymptotically stable, endemic equilibrium. We derive the respective fitness gradients, 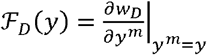 and 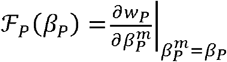, from equations (6)-(7) (omitted for brevity) and find singular strategies, *y*^*^ and 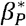, by numerically solving ℱ_*D*_ (*y*\*)= 0 and 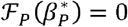. Singular strategies are evolutionarily stable if 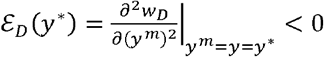 and 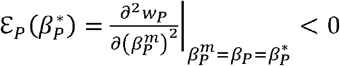, respectively. For parasite evolution only, we determine convergence stability by numerically evaluating the derivative 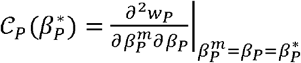 and checking that 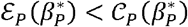. In the case of coevolution, we assume equal mutation rates for defensive symbionts and parasites, and determine strong convergence stability using the method presented in (Leimer 2009) (see *Supplementary Material*).

We assume that the defensive symbiont is introduced into a well-established host-parasite system, with the parasite at its unique continuously stable strategy (CSS) in the absence of the defensive symbiont (see *Supplementary Material*), 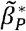, which is given by

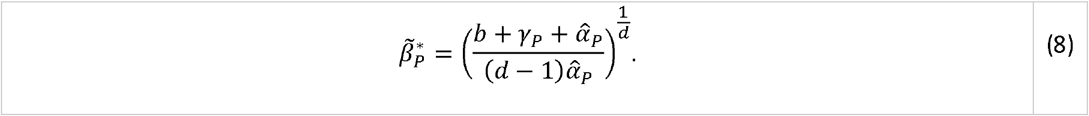

In addition to exploring the effects of the defensive symbiont on the (co)evolution of virulence and host protection, we measure the net effect on the host population size and change in the average host mortality rate (relative to the initial symbiont-free population). The net effect on the host population size is measured by comparing the steady state in the presence and absence of the defensive symbiont, *N*\* (*y,β*_*p*_) and *Ñ*^*^ respectively. Similarly, we calculate the average disease-associated mortality rate at equilibrium in the presence and absence of the defensive symbiont, *r*^*^ *y,β*_*p*_) and 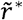, respectively as:

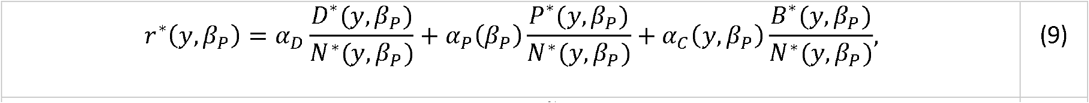

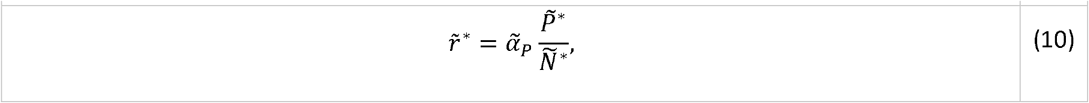

where we have explicitly written the dependence of the trait variables on the steady state values. Note that the values for *Ñ*^*^ and 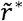 are constants calculated at the singular strategy for a parasite circulating in the absence of the defensive symbiont, where we begin all of our evolutionary simulations. We then define the following two measures to determine the net effects on the host population following the introduction of the defensive symbiont:

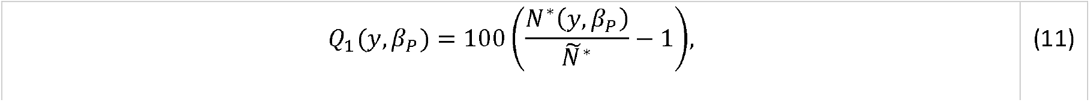

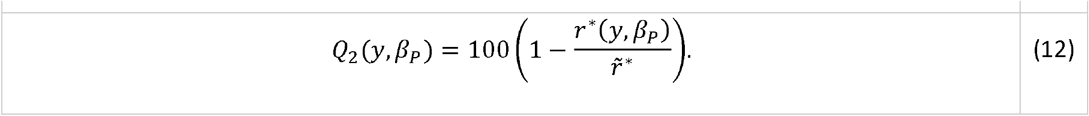

The first measure (equation 11) is the percentage increase in the host population size and the second measure (equation 12) is the percentage decrease in the disease-associated mortality rate.

### Simulations

The above analysis makes two key assumptions: (1) that there is a separation of the ecological and evolutionary time scales (i.e. mutations are rare), and (2) that selection is weak, so that mutations only have a small phenotypic effect (i.e. traits are continuous). We relax these assumptions in our simulations by allowing new mutants to arise before the ecological dynamics are close to their ecological attractor and by discretising the trait space so that new mutations have small but finite effects. Simulations proceed as follows (described for the coevolution case). We initialise a resident population which has a defensive symbiont protection level of *y*^*r*^ and a parasite transmission of 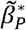 as defined in equation (8). We simulate the ecological dynamics (1)-(4) for a total (arbitrary) time of *T*_*eco*_ = 100. We choose either the defensive symbiont or parasite population with equal probability and introduce a mutant at low frequency with trait value differing from the resident by a small amount, *∈* _*D*_ or *∈* _*p*_. We then run the ecological dynamics again for another *T*_*eco*_ time units, remove any phenotypes that have dropped below a frequency of *ε* =10^−4^ (this threshold is arbitrary) and then introduce a new mutant again, by firstly choosing the defensive symbiont or parasite with equal probability and then choosing a trait to mutate proportional to its frequency. This continues for a total of *T*_*evo*_ evolutionary time-steps.

## Results

We begin by exploring how the introduction of a (non-evolving) defensive symbiont affects the quantitative and qualitative evolution of parasite virulence, before considering coevolution of both microbes.

### Defensive symbionts that confer tolerance always select for increased virulence

The introduction of a non-evolving defensive symbiont, which confers a fixed level of tolerance to parasitised hosts, always leads to selection for higher parasite virulence (Fig. 2a). This is because the defensive symbiont not only directly reduces virulence when present with the parasite (hence, reducing the cost to the parasite of elevated virulence), but also competes with the parasite for hosts (thus increasing selection for a higher transmission rate, and hence higher virulence) even when providing little to no protection. The latter effect is more subtle and is typically weaker but is evident when the defensive symbiont confers no protection to the host (*y* = 0), as the parasite still evolves increased virulence due to increased competition for hosts. The strength of the first effect depends on both the level (*y*) and cost (*c*_1_) of conferred protection, which together determine how often the parasite shares a host with a symbiont (Fig. 2c). When the cost to the defensive symbiont of conferring tolerance (*c*_1_) is sufficiently low, greater host protection (*y*) always selects for higher parasite virulence because the parasite frequently shares hosts with the defensive symbiont, and so benefits from decreased realised virulence due to tolerance conferred to the host by the symbiont. However, when the cost of host protection is relatively high, fewer hosts harbour the defensive symbiont and so the parasite is less likely to benefit from conferred tolerance, resulting in evolved virulence peaking at an intermediate level of host protection (Fig. 2a).

**Fig. 2:**
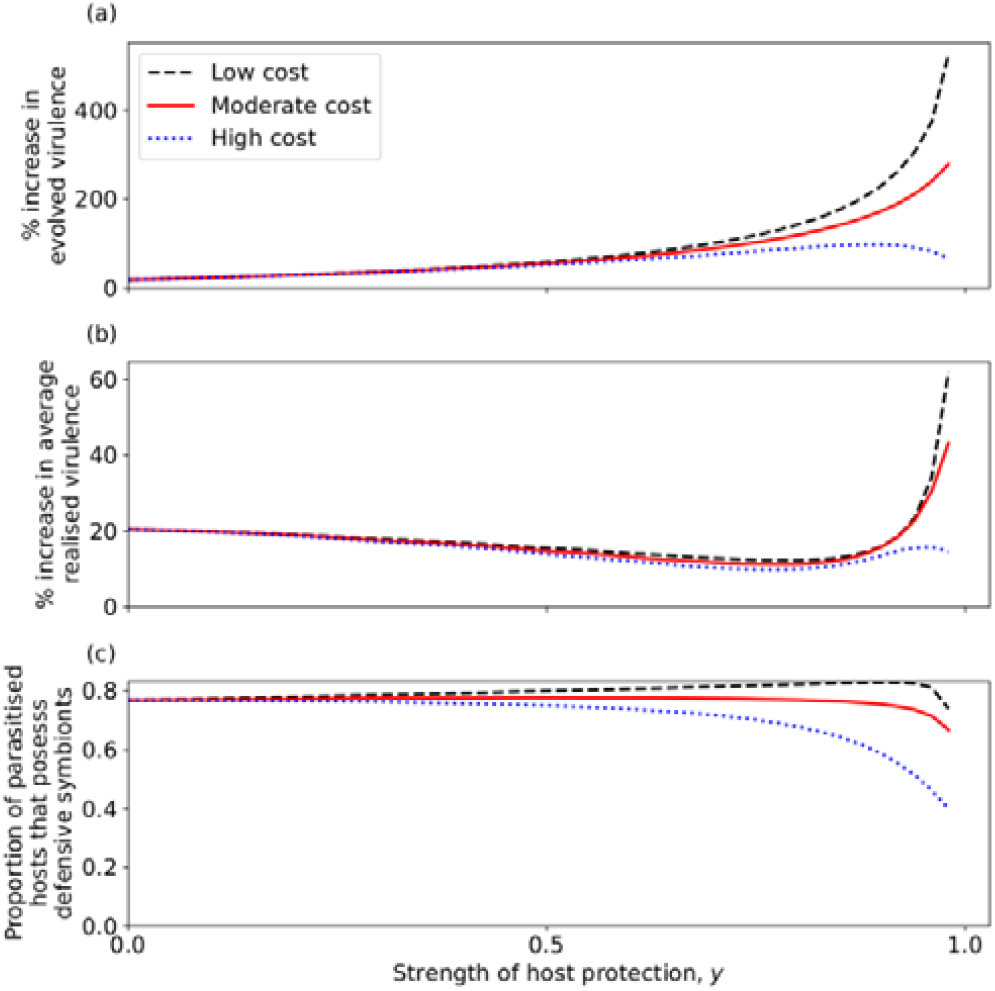
Evolution of parasite virulence following the introduction of the defensive symbiont. (a) Evolved virulence relative to the initial stable level of virulence in the absence of the defensive symbiont. (b) The percentage increase in average realised virulence compared with the absence of the defensive symbiont. (c) The proportion of parasitised hosts that possess defensive symbionts. The black dashed line corresponds to relatively low costs to the defensive symbiont of conferring protection, the red solid line to moderate costs and the blue dotted line to relatively high costs. Values of strength of cost have been chosen to represent strengths of cost across the entire range. All other parameters are as in Table 1.

As the defensive symbiont confers tolerance to the host, higher evolved virulence does not necessarily imply that realised virulence will be higher. Yet, following the introduction of the defensive symbiont, there is always an increase in average realised virulence (i.e. the average level of virulence experienced by parasitised hosts, with or without the defensive symbiont; Fig. 2b). Average realised virulence is markedly lower than the increase in evolved virulence (Fig. 2a-b) due to the presence of the defensive symbiont, but hosts that do not possess the defensive symbiont will experience the full increase in virulence. Average realised virulence is minimised at an intermediate level of host protection, where there are relatively more hosts harbouring both microbes (Fig. 2c), and at high levels of protection there can be a sharp increase in average realised virulence due to a combination of strong selection for virulence (Fig. 2a) and fewer hosts possessing the defensive symbiont (Fig. 2c).

**Table 1:** Default parameter values for the model (1)-(4).

| Parameter | Description | Default value |
| --- | --- | --- |
| $a$ | Maximum per-capita host birth rate | 1.0 |
| $b$ | Host natural mortality rate | 0.25 |
| $c_1$ | Defensive symbiont cost strength parameter | 0.25 |
| $c_2$ | Defensive symbiont cost shape parameter | 2 |
| $d$ | Power-law for parasite virulence cost | 2 |
| $q$ | Strength of density-dependent competition on host reproduction | 0.25 |
| $T_{\text{eco}}$ | Duration for ecological time steps | 100 |
| $T_{\text{evo}}$ | Duration for evolutionary simulations | 2000 |
| $\alpha_D$ | Cost of harbouring the defensive symbiont | 0.1 |
| $\hat{\alpha}_P$ | Virulence of a parasite that cannot transmit | 0.1 |
| $\hat{\beta}_D$ | Transmission rate of a defensive symbiont with no cost of protection | 2 |
| $\gamma_D$ | Host recovery rate for defensive symbiont | 0.05 |
| $\gamma_P$ | Host recovery rate for parasite | 0.05 |
| $\varepsilon$ | Extinction threshold | $10^{-4}$ |

### Defensive symbionts can drive parasite diversification

In addition to selecting for higher parasite virulence, the defensive symbiont can also drive diversification when tolerance is maximised or very close to being maximised (*y* ≈ 1), causing the parasite to branch into two subpopulations (Fig. 3). One of these subpopulations has a high level of virulence (and transmission), and is primarily found in hosts that also harbour the defensive symbiont, while the other evolves a much lower level of virulence and is primarily found in hosts that do not harbour the defensive symbiont (Fig. 3bii). Note that when tolerance is maximised at *y* = 1, parasite virulence is completely negated in hosts that possess defensive symbionts, but the two strains are maintained in the population due to their contrasting strategies in isolation (infecting fully susceptible hosts compared to those that harbour the defensive symbiont) and the frequency with which they co-occur with the defensive symbiont. Evolutionary branching in parasite virulence occurs when the strength of the cost to the defensive symbiont is within a relatively narrow range. When the costs of host protection are below this range, there is only runaway selection for virulence (Fig. 3ai), and when the costs are above this range, there may be runaway selection for virulence or a stable level of virulence may evolve (Fig. 3aiii).

**Fig. 3:**
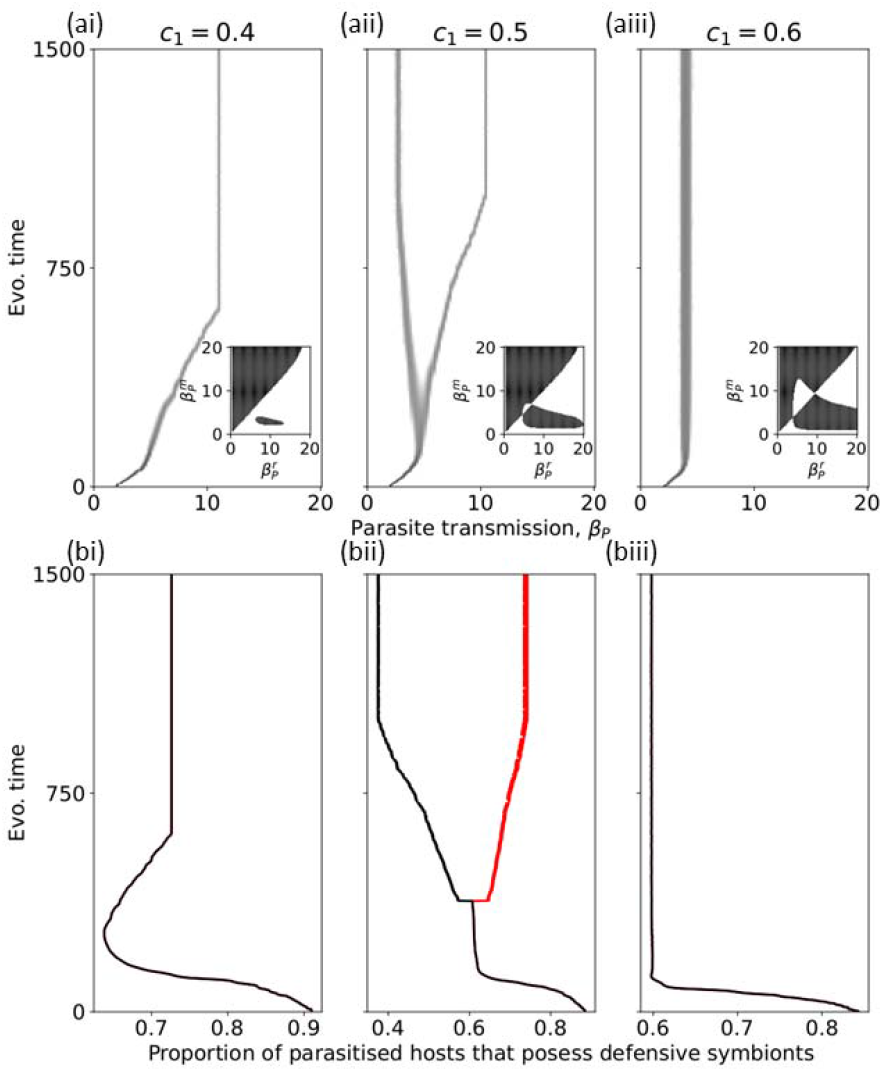
Parasite diversification driven by a defensive symbiont. Row a: evolutionary trajectories of parasite transmission with inset pairwise invasion plots (PIPs). Black regions in the PIPs show where the mutant can invade (where 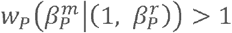), and white regions are where it cannot. Row b: the proportion of parasitised hosts which also possess the defensive symbiont. For bii, the red line (right branch) corresponds to the high virulence strain, and the black line (left branch) corresponds to the low virulence strain. Costs of host protection: (column i) c_1_ =0.4, (column ii) c_1_ =0.5 and (column iii) c_1_ =0.6. These values of cost strength represent the narrow range of values with a qualitative change in evolutionary behaviour. All other parameters as in Table 1.

### Symbiont-parasite coevolution can be detrimental to the host population

We now allow the level of protection conferred by the defensive symbiont to coevolve with parasite virulence. The parasite, as before, is initialised to its stable level of virulence (equation 8) in the absence of the defensive symbiont. We then introduce the defensive symbiont at different initial levels of protection to determine if coevolution results in (i) increased or decreased conferred protection and (ii) a net cost or benefit to the host population.

We first determine the range of possible evolutionary outcomes for the defensive symbiont as the cost parameters associated with host protection vary (Fig. 4). It is well-established that trade-off shapes determine qualitative evolutionary outcomes (Andrew Hoyle et al. 2008) and the range of outcomes in our model and when they occur is consistent with previous theory (Ashby and King 2017). Under decelerating trade-offs (*c*_2_ < 0), the defensive symbiont either maximises or minimises host protection (potentially depending on the initial level of protection; Fig. 4), as a small increase from no protection (*y* = 0) is relatively costly, whereas changes at higher levels of protection are less costly. The defensive symbiont therefore either overcomes the initial cost and experiences runaway selection for maximal protection, or experiences selection against host protection. When the costs of host protection accelerate (*c*_2_ > 0), the defensive symbiont maximises protection if the strength of the cost is sufficiently low, and evolves to either an intermediate level of protection or no protection if the strength of the cost is sufficiently high (Fig. 4).

**Fig. 4:**
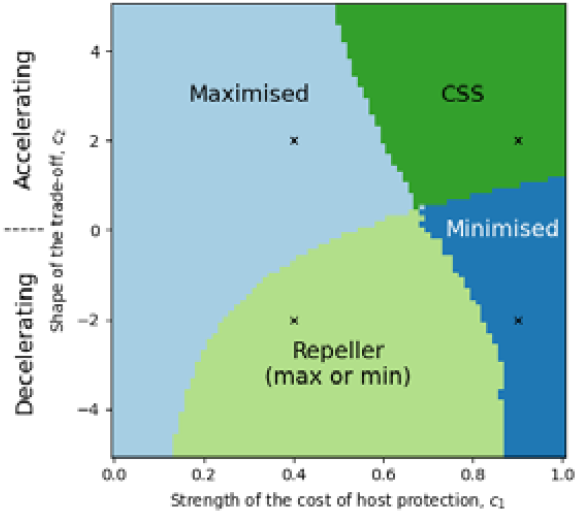
Classification of the coevolutionary outcome for the defensive symbiont as a function of the two cost function parameters;, the strength of cost ranging from 0 (no cost) to 1 (maximal cost), and the shape of the trade-off with transmission: accelerating, linear, decelerating. The repeller region results in the defensive symbiont either maximising or minimising host protection depending on the initial level of protection. The CSS region corresponds to a continuously stable strategy at an intermediate level of protection. The parasite is also evolving, but its evolutionary outcome is not shown as it always tends to a CSS. Black crosses correspond to the 4 parameter pairs used in Fig. 5 to demonstrate the different qualitative behaviours. All other parameters as in Table 1.

We now consider how virulence coevolves with host protection to determine the net effect on the host population following the introduction of the defensive symbiont (Fig. 5). First, we find that while a defensive symbiont may initially increase the host population size, the host appears eventually to always suffer a decrease in population size due to parasite-symbiont coevolution, regardless of the initial strength of protection (indicated by the terminus of each evolutionary trajectory residing in regions with a negative percentage increase in host population size). This seems to occur for one or more of the following three reasons: (1) the symbiont may experience selection against tolerance, resulting in a reduction or even loss of host protection; (2) the defensive symbiont incurs a small cost to the host; and (3) while the defensive symbiont may confer tolerance to some hosts, the parasite subsequently experiences selection for higher virulence, and so hosts without the defensive symbiont experience higher virulence.

**Fig. 5:**
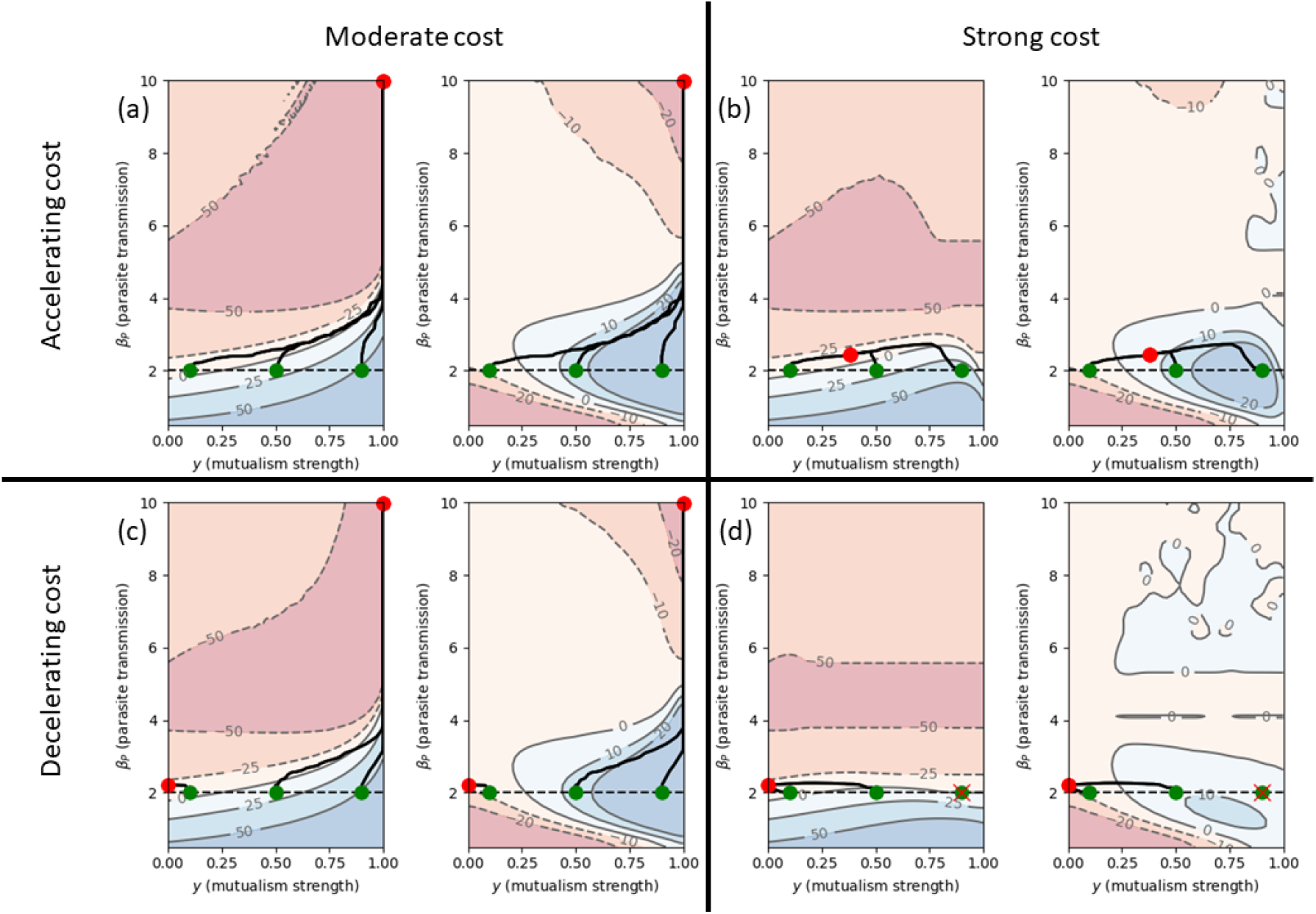
Heatmaps for the changes in population size and death rate (as given in equations (11)-(12)) for various mutualist cost functions. We show moderate cost (c_1_ =0.4) in the left column and strong cost (c_1_ =0.9) in the right, with accelerating cost (c_2_ =2) and decelerating cost (c_2_ =−2) in the top and bottom rows respectively. Colours and values on the contour plots denote percentage changes for a given trait space pair (y,β_P_). Green dots are the initial value, solid black lines denote an evolutionary trajectory in trait space, and the red dots are the ends. The black dashed line is the CSS value for the parasite transmission when it is the only microbe in circulation. The values of c_1_ and c_2_ have been chosen to represent the four qualitative behaviours for the defensive symbiont that we have seen in Fig. 4.

Although there is always eventually a net-negative effect on the host population size following parasite-symbiont coevolution, the same is not necessarily true for realised virulence (i.e. the average disease-associated mortality rate). In many cases an initially positive effect on average realised virulence is followed by a long-term negative effect (as observed for the host population size measure above), but when the costs of protection are sufficiently strong and accelerate, there is a reduction in average realised virulence (Fig. 5b).

When the costs of protection accelerate, the parasite and symbiont coevolve to co-continuously stable strategies (Fig. 5a and b), but when the costs of protection decelerate, the outcome may depend on the initial conditions, with sufficiently low levels of initial protection leading to selection against any protection and a minor increase in parasite virulence (Fig. 5c and d), and sufficiently high levels of initial protection leading to selection for maximal protection and high virulence (Fig. 5c). Somewhat paradoxically, this means that the introduction of a highly protective symbiont can lead to a much larger negative effect on the host population than the introduction of a symbiont that confers only a low level of protection.

## Discussion

Defensive symbionts are found throughout the natural world and are potentially important agents of biocontrol, yet the robustness of host protection and their eco-evolutionary impacts on parasite evolution are poorly understood. In this study, we have theoretically explored the (co)evolutionary dynamics of parasite virulence and host protection by a defensive symbiont in the form of tolerance. We have investigated the behaviour of both the parasite and the defensive symbiont, as well as the net effect on the host population. We have shown that the parasite will always evolve to be more virulent following the introduction of a tolerance-conferring defensive symbiont, and (for every parameter combination tested) this always has a negative impact on the host population size even if the defensive symbiont evolves to confer maximum host protection. Furthermore, our model reveals that the defensive symbiont can cause diversification in the parasite population for sufficiently high levels of host protection, leading to the coexistence of low and high virulence phenotypes. Overall, our results suggest that the introduction of tolerance-conferring defensive symbionts is likely to lead to higher evolved and realised virulence, resulting in a net negative impact on the host population.

Higher virulence always evolves in our model because the defensive symbiont confers protection to the host by ameliorating the disease-associated mortality rate, which increases the average infectious period in coinfected hosts – those harbouring the defensive symbiont and parasite simultaneously. Although more virulent parasites experience a sub-optimal level of virulence in hosts that do not harbour the defensive symbiont, this is more than offset by having a higher transmission rate in coinfections. Thus, the prevalence of the defensive symbiont, and hence the frequency of coinfections, plays a crucial role in determining the strength of selection for increased virulence. The fact that tolerance-conferring symbionts always select for higher virulence mirrors the literature on imperfect vaccination. Gandon et al. (2001) showed theoretically how partially effective vaccines that prevent or reduce disease (i.e., confer tolerance) but do not prevent transmission select for higher virulence, a prediction that has since been confirmed for Marek’s disease in chickens (Read et al. 2015). Imperfectly vaccinated individuals are analogous to hosts who harbour the defensive symbiont in our model; in both cases, the host experiences lower virulence while still being able to transmit the infectious agent, weakening the evolutionary trade-off between transmission and virulence and shifting the balance of selection towards higher virulence. Note that in our model there is no explicit reduction in the pathogen growth rate due to tolerance (although this need not be the case in general) and there is no direct effect on the transmission rate. Instead, the defensive symbiont confers protection by reducing the negative effects of infection (akin to the “anti-toxin” resistance in Gandon et al. (2001)). While we are not aware of any experimental studies that have explored the evolution of virulence in the presence of a tolerance-conferring symbiont, the strong parallels with imperfect vaccination suggest that such symbionts should indeed select for higher parasite virulence.

Although we found that the introduction of a tolerance-conferring defensive symbiont always selects for higher virulence, this is not necessarily the case for all symbionts. For example, Nelson and May (2017, 2020) have shown that when the shared costs of virulence are sufficiently low, communities of symbionts remain mutualistic and evolve lower virulence, and that in certain cases pathogens can evolve towards mutualism in these communities. There are several key differences between our model and those studied by Nelson and May (2017, 2020). In particular, Nelson and May consider shared (additive) costs of virulence, whereas in our model host protection directly (i.e. multiplicatively) reduces the virulence of the parasite. Furthermore, the mutualistic effects of our defensive symbiont are context-dependent, as the benefits of protection are only realised in infected hosts, whereas the mutualistic effects observed by Nelson and May occur regardless of whether other symbionts are present. Our model therefore highlights the importance of context-dependent mutualisms and direct interactions between mutualists and parasites for the evolution of virulence.

Even if evolved virulence is higher in the presence of the defensive symbiont, the realised virulence experienced by hosts with the symbiont can be lower due to host protection. However, hosts without the defensive symbiont will experience increased virulence, and so the frequency of coinfections will determine the variance in the realised level of virulence experienced by parasitised hosts. While the net effect of the defensive symbiont on the host population size might initially be positive, we have shown that this is not evolutionary robust, either due to selection for higher parasite virulence (even if selection also favours higher host protection by the defensive symbiont, as in Fig. 5a), or due to selection against host protection (Fig. 5c-d). However, if the goal is to reduce the average virulence experienced by infected hosts rather than to maximise host population size, then it is possible to achieve modest gains in host survival provided the cost of conferring host protection accelerates with greater host protection and the overall strength of costs are sufficiently high (Fig. 5b).

Our results have critical implications for the use of defensive symbionts as biocontrol agents, with tolerance-conferring symbionts likely to be a poor choice for long-term infectious disease control at the population level. Moreover, our model demonstrates the need to investigate the possible evolutionary dynamics of both defensive symbionts and parasites when considering the use of biocontrols, as short-term ecological dynamics may be a poor predictor of long-term outcomes. Counter-intuitively, our model reveals that under certain trade-offs (when costs of host protection have increasing returns, i.e. are decelerating), the introduction of a more protective defensive symbiont can lead to far worse outcomes for the host population in the long-term compared than if a less protective symbiont is introduced (Fig. 5c). Decelerating trade-offs often produce evolutionary repellers (e.g., Ashby and King 2017) as they impose a high initial cost when investment is low (here, this selects against protection and leads to little change in virulence if the initial level of protection is low), and a relatively low additional cost when investment is high (here, this selects for higher protection, and in turn, higher virulence, when the initial level of protection is sufficiently high). Due to the complex nature of eco-evolutionary feedbacks in these systems and the potential for unexpected evolutionary trajectories, we therefore urge caution in the use of tolerance-conferring symbionts.

Our final key result is that the defensive symbiont can drive parasite diversification into high and low virulence phenotypes. This occurs because the defensive symbiont adds an additional feedback on the parasite population, which allows the different phenotypes to specialise on hosts that either lack or possess the defensive symbiont. However, we found that the level of tolerance conferred by the symbiont must be very high for diversification to occur, which suggests that although this is theoretically possible, it is unlikely to be common in real populations. Nevertheless, the fact that a defensive symbiont can facilitate parasite diversification emphasises the importance of considering community effects on host and parasite diversity, and this finding follows a general pattern in recent theoretical studies where the addition of a third species induces diversification in the host or parasite (Best 2018; Kisdi, Geritz, and Boldin 2013; Wood and Ashby 2023; Hoyle, Best, and Bowers 2012). For example, the addition of a predator that differentially feeds on infected hosts has been shown to lead to diversification in host resistance (Hoyle, Best, and Bowers 2012) and parasite virulence (Kisdi, Geritz, and Boldin 2013; Best 2018). Similarly, the introduction of a hyperparasite (a parasite of the parasite) can also cause diversification in parasite virulence (Wood and Ashby 2023), which potentially presents problems for biocontrol, as with tolerance-conferring symbionts.

To date, few studies have experimentally explored the evolution of parasite virulence in the presence of defensive symbionts (King et al. 2016; Ford et al. 2017; G. May et al. 2022). May et al. (2022) have shown that when a plant host (*Zea mays*) is infected by a pathogenic fungus (*Ustilago maydis*), parasite fitness is maximised at higher levels of virulence in the presence of a defensive symbiont (*Fusarium verticillioides*), in agreement with our model. However, Ford et al. (2016) experimentally coevolved pathogenic *Staphylococcus aureus* and protective *Enterococcus faecalis* in *Caenorhabditis elegans* hosts, which led to a reduction in pathogen virulence; the same study system has also been shown to generate fluctuating selection dynamics in pathogen virulence and host protection (Ford et al. 2017). The contrast with our results is because *E. faecalis* confers protection through the production of antimicrobial superoxides which directly inhibit *S. aureus* rather than conferring tolerance to the host. The stark contrast in evolutionary outcomes with our model emphasises the importance of understanding the mechanism of host protection. Moreover, most studies of defensive symbionts focus on those that confer protection in terms of a reduced parasite load (e.g., due to interference competition) (Hoang and King 2022), and tolerance-conferring symbionts are understudied. Indeed, we are aware of only one study that explicitly shows defensive symbionts conferring tolerance to the host, with *Bacteroides fragilis* conferring tolerance by inducing the production of anti-inflammatory proteins against an experimental colitis caused by the bacterium *Heliobacter hepaticus* (Mazmanian, Round, and Kasper 2008).

Given that many defensive symbionts confer host protection through other mechanisms (including through upregulation of host immune responses (Ford, Drew, and King 2022), the eco-evolutionary implications of introducing different types of symbionts should be explored in future theoretical studies. Furthermore, we have implicitly modelled how the defensive symbiont and parasite interact at the within-host level, along with trade-offs between transmission and tolerance or virulence. This simplification makes the model much more tractable, but an important direction for future research is to explicitly model the within-host dynamics and couple these to between-host transmission. Coupling within- and between-host modelling has been shown to provide new insights than population-level modelling on its own cannot provide. Modelling at each level explicitly means that we can investigate conflicting selection, i.e. where the most successful phenotype at one level is not necessarily the most successful at the other (Frank 1996; van Baalen and Sabelis 1995; B. R. Levin and Bull 1994). The overall evolutionary outcomes will heavily depend on this conflicting selection that cannot be fully captured by simply modelling at the between-host level only. Another important consequence of explicit within-host modelling is providing insights into trade-offs and recovery rates. Several nested models have shown that any trade-offs depend heavily on the within-host dynamics (Ganusov, Bergstrom, and Antia 2002; Gilchrist and Sasaki 2002; André, Ferdy, and Godelle 2003; Alizon and van Baalen 2005; Gilchrist and Coombs 2006), whilst recovery rates, which are traditionally considered to be constant and independent of other parameters, have been shown to be important for the evolution of virulence (Ganusov and Antia 2006; André and Gandon 2006). In the context of defensive symbiont-parasite dynamics, explicit within-host modelling will allow for a greater understanding of how a range of different mechanisms, such as interference competition, resource competition, spite, priority effects and interactions with the host immune system impact on the evolution of virulence and host protection.

Overall, our model reveals how tolerance-conferring defensive symbionts typically have a net negative impact on the host population over the long-term as they always select for higher parasite virulence and are therefore poor candidates for biocontrol.

## Acknowledgements

BA is funded by Natural Environment Research Council grants NE/N014979/1 and NE/V003909/1. CAS is funded by Natural Environment Research Council grant NE/V003909/1. We acknowledge the support of the Natural Sciences and Engineering Research Council of Canada (NSERC). Nous remercions le Conseil de recherches en sciences naturelles et en génie du Canada (CRSNG) de son soutien. PIPPS receives funding from the BC Ministry of Health.

## Data accessibility

The Python code for the implementation of this model can be found here: https://github.com/CameronSmith50/Defensive-Symbiosis

