## Supplementary Information for "Tolerance-conferring defensive symbionts and the evolution of parasite virulence"

**Abstract**

Defensive symbionts in the host microbiome can confer protection from infection or reduce the harms of being infected by a parasite. Defensive symbionts are therefore promising agents of biocontrol that could be used to control or ameliorate the impact of infectious diseases. Previous theory has shown how symbionts can evolve along the parasitism-mutualism continuum to confer greater or lesser protection to their hosts, and in turn how hosts may coevolve with their symbionts to potentially form a mutualistic relationship. However, the consequences of introducing a defensive symbiont for parasite evolution and how the symbiont may coevolve with the parasite have yet to be explored theoretically. Here, we investigate the ecological and evolutionary implications of introducing a tolerance-conferring defensive symbiont into an established host-parasite system. We show that while the defensive symbiont may initially have a positive impact on the host population, parasite and symbiont evolution tend to have a net negative effect on the host population in the long-term. This is because the introduction of the defensive symbiont always selects for an increase in parasite virulence, and may cause diversification into high- and low-virulence strains. Even if the symbiont experiences selection for greater host protection, this simply increases selection for virulence in the parasite, resulting in a net negative effect on the host population. Our results therefore suggest that tolerance-conferring defensive symbionts may be poor biocontrol agents for population-level infectious disease control.

### 1 Calculation of fitness functions and selection gradients

In this section, we will derive the fitness functions and selection gradients given in the main text. We will show the full calculation for the mutualist, but the calculation for the parasite is similar.

Recall from the main text, the dynamics (1)-(4) that defines a resident population of mutualists and parasites circulating through a host population. We re-write this here, writing the trait values with a superscript "$r$" to denote they are resident traits:

$$\frac{\mathrm{d}H}{\mathrm{d}t} = \nu(N) - \left[b + \beta_D(y^r)(D+B) + \beta_P^r(P+B)\right] H + \gamma_D D + \gamma_P P, \tag{S1}$$

$$\frac{\mathrm{d}D}{\mathrm{d}t} = \beta_D(y^r)H(D+B) - \left[b + \gamma_D + \alpha_D + \beta_P^r(P+B)\right] D + \gamma_P B, \tag{S2}$$

$$\frac{\mathrm{d}P}{\mathrm{d}t} = \beta_P^r H(P+B) - \left[b + \gamma_P + \alpha_P(\beta_P^r) + \beta_D(y^r)(D+B)\right] P + \gamma_D B, \tag{S3}$$

$$\frac{\mathrm{d}B}{\mathrm{d}t} = \beta_D(y^r)P(D+B) + \beta_P^r D(P+B) - \left[b + \gamma_D + \gamma_P + \alpha_B(y^r, \beta_P^r)\right] B, \tag{S4}$$

with all parameters and state variables as described in the main text. We assume that we introduce a mutant mutualist with trait value $y^m$ into a resident population which has reached equilibrium. We denote by $X^*$ for $X \in \{H, D, P, B, N\}$ to be the steady states of the resident population. Then the ODEs for the mutant mutualist, $D^m$, and those coinfected by the mutant mutualist and resident parasite, $B^m$, is given by:

$$\frac{\mathrm{d}D^m}{\mathrm{d}t} = \beta_D(y^m)H^*(D^m + B^m) - \left[b + \gamma_D + \alpha_D + \beta_P^r(P^* + B^*)\right] D^m + \gamma_P B^m, \tag{S5}$$

$$\frac{\mathrm{d}B^m}{\mathrm{d}t} = \beta_D(y^m)P^*(D^m + B^m) + \beta_P^r D^m(P^* + B^*) - \left[b + \gamma_D + \gamma_P + \alpha_B(y^m, \beta_P^r)\right] B^m. \tag{S6}$$

We will use the next generation method (Diekmann et al., 2010) in order to calculate an expression which is sign equivalent to the fitness of the rare mutant.

We decompose the Jacobian matrix of first derivatives into two components, $J = F + V$, where $F$ describes the creation of new mutant infections and $V$ contains all other terms:

$$\begin{aligned} J &= \begin{bmatrix} \beta_D(y^m)H^* - \left[b + \gamma_D + \alpha_D + \beta_P^r(P^* + B^*)\right] & \beta_D(y^m)H^* + \gamma_P \\ \beta_D(y^m)P^* + \beta_P^r(P^* + B^*) & \beta_D(y^m)P^* - \left[b + \gamma_D + \gamma_P + \alpha_B(y^m, \beta_P^r)\right] \end{bmatrix} \\ &= \beta_D(y^m)\begin{bmatrix} H^* & H^* \\ P^* & P^* \end{bmatrix} + \begin{bmatrix} -\left[b + \gamma_D + \alpha_D + \beta_P^r(P^* + B^*)\right] & \gamma_P \\ \beta_P^r(P^* + B^*) & -\left[b + \gamma_D + \gamma_P + \alpha_B(y^m, \beta_P^r)\right] \end{bmatrix} \\ &=: F + V. \end{aligned}$$

This decomposition allows us to calculate the next generation matrix, $N_G = -FV^{-1}$. We can then find our quantity of question by calculating the leading eigenvalue of this matrix:

$$N_G = \frac{\beta_D(y^m)}{\det(V)}\begin{bmatrix} H^*\left[b + \gamma_D + \gamma_P + \alpha_B(y^m, \beta_P^r) + \beta_P^r(P^* + B^*)\right] & H^*\left[b + \gamma_D + \gamma_P + \alpha_D + \beta_P^r(P^* + B^*)\right] \\ P^*\left[b + \gamma_D + \gamma_P + \alpha_B(y^m, \beta_P^r) + \beta_P^r(P^* + B^*)\right] & P^*\left[b + \gamma_D + \gamma_P + \alpha_D + \beta_P^r(P^* + B^*)\right] \end{bmatrix},$$

where

$$\det(V) = \left[b + \gamma_D + \gamma_P + \alpha_B(y^m, \beta_P^r)\right]\left[b + \gamma_D + \alpha_D + \beta_P^r(P^* + B^*)\right] - \gamma_P \beta_P^r(P^* + B^*).$$

This matrix has two eigenvalues, one which is zero and then the leading eigenvalue:

$$w_M(y^m|y^r, \beta_P^r) = \frac{\beta_D(y^m)A_M(y^m|y^r, \beta_P^r)}{B_M(y^m|y^r, \beta_P^r)}, \tag{S7}$$

where

$$\begin{aligned} A_M(y^m|y^r, \beta_P^r) = H^* \left[b + \gamma_D + \gamma_P + \alpha_B(y^m, \beta_P^r) + \beta_P^r(P^* + B^*)\right] \\ + P^* \left[b + \gamma_D + \gamma_P + \alpha_D + \beta_P^r(P^* + B^*)\right], \end{aligned} \tag{S8}$$

$$B_M(y^m|y^r, \beta_P^r) = \det(V). \tag{S9}$$

If the fitness function is above 1, the mutant is able to invade into the population, while if this value is below 1, it is not. Note also that if $y^m = y^r$, then $w_M(y^m|y^r, \beta_P^r) = 1$ because there is no fitness advantage or disadvantage for the mutant due to it being the same as the resident.

We now calculate the selection gradient, $H_M(y, \beta_P)$. This is defined as the first derivative of the fitness function with respect to the mutant trait and evaluated at $y^r = y^m = y$ and $\beta_P = \beta_P^r$. We write

$$\begin{aligned} s_M(y, \beta_P) &= \frac{\partial w_M}{\partial y^m}(y^m|y^r, \beta_P^r)\bigg|_{\substack{y^r = y^m = y,\\ \beta_P^r = \beta_P}} \\ &= \frac{1}{B_M(y|y, \beta_P)}\left[\frac{\mathrm{d}\beta_D}{\mathrm{d}y^m}(y^m)A_M(y^m|y^r, \beta_P^r) + \beta_D(y^m)\frac{\partial A_M}{\partial y^m}(y^m|y^r\beta_P^r)\right. \\ &\qquad \left.\left. - \frac{\beta_D(y^m)A_M(y^m|y^r, \beta_P^r)}{B_M(y^m|y^r, \beta_P^r)}\frac{\partial B_M}{\partial y^m}(y^m|y^r\beta_P^r)\right]\right|_{\substack{y^r = y^m = y,\\ \beta_P^r = \beta_P}} \\ &= \frac{1}{B_M(y|y, \beta_P)}\left[\frac{\mathrm{d}\beta_D}{\mathrm{d}y^m}(y^m)A_M(y^m|y^r, \beta_P^r) + \beta_D(y^m)\frac{\partial A_M}{\partial y^m}(y^m|y^r\beta_P^r)\right. \\ &\qquad \left.\left. - w_M(y^m|y^r, \beta_P^r)\frac{\partial B_M}{\partial y^m}(y^m|y^r, \beta_P^r)\right]\right|_{\substack{y^r = y^m = y,\\ \beta_P^r = \beta_P}}. \end{aligned}$$

Finally, we calculate each of these derivatives that we require. Firstly, the derivative of $\beta_D$:

$$\begin{aligned} \frac{\mathrm{d}\beta_D}{\mathrm{d}y^m}(y^m) &= \frac{\mathrm{d}}{\mathrm{d}y^m}\left(\tilde{\beta}_M(1 - c(y_m))\right) \\ &= \tilde{\beta}_M\left(1 - \frac{\mathrm{d}c}{\mathrm{d}y_m}(y_m)\right) \\ &= \tilde{\beta}_M\left(1 - \frac{c_1 c_2 \exp\{c_2 x\}}{1 - \exp\{c_2\}}\right). \end{aligned}$$

Next, we differentiate the $A_M$ function:

$$\begin{aligned} \frac{\partial A_M}{\partial y^m}(y^m|y^r, \beta_P^r) &= H^*\frac{\partial \alpha_B(y^m, \beta_P^r)}{\partial y^m} \\ &= H^*\frac{\partial}{\partial y^m}\left(\alpha_D + (1 - y^m)\alpha_P(\beta_P^r))\right) \\ &= -H^*\alpha_P(\beta_P^r). \end{aligned}$$

Finally, the derivative for the $B_M$ function:

$$\begin{aligned} \frac{\partial B_M}{\partial y^m}(y^m|y^r\beta_P^r) &= \frac{\partial \alpha_B}{\partial y^m}(y^m, \beta_P^r)\left[b + \gamma_D + \alpha_D + \beta_P^r(P^* + B^*)\right] \\ &= -\alpha_P(\beta_P^r)\left[b + \gamma_D + \alpha_D + \beta_P^r(P^* + B^*)\right]. \end{aligned}$$

#### 2 Parasite only system

Throughout the manuscript, we initialise our simulations assuming that the parasite has been circulating on its own long enough to reach evolutionary convergence. Here, we describe the parasite-only system, it's fitness function and calculate the transmission value to which it converges.

We begin by defining the parasite-only system, where throughout this section we will indicate state variables with a tilde ($\sim$) in order to distinguish from the full system. The ODE system is given by:

$$\frac{\mathrm{d}\tilde{H}}{\mathrm{d}t} = \nu(\tilde{N}) - \left[b + \tilde{\beta}_P \tilde{P}\right] + \gamma_P \tilde{P}, \tag{S10}$$

$$\frac{\mathrm{d}\tilde{P}}{\mathrm{d}t} = \left[\tilde{\beta}_P \tilde{H} - b - \gamma_P - \alpha_P(\tilde{\beta}_P)\right] \tilde{P}, \tag{S11}$$

where all notation is the same as in the main text. This ODE system has two endemic steady states, which are given by the solution of:

$$\tilde{H}^* = \frac{b + \gamma_P + \alpha_P(\tilde{\beta}_P)}{\tilde{\beta}_P}, \tag{S12}$$

$$0 = \left(\tilde{P}^*\right)^2 - \frac{a - 2q\tilde{H}^* - \tilde{\beta}_P \tilde{H}^* + \gamma_P}{q}\tilde{P}^* - \frac{\tilde{H}^*(a - b - q\tilde{H}^*)}{q}. \tag{S13}$$

We now write the fitness function and associated selection gradient for this parasite-only system for a mutant strain with transmission $\tilde{\beta}_P^m$ introduced into a population containing a resident strain with transmission $\tilde{\beta}_P^r$ as:

$$\tilde{w}(\tilde{\beta}_P^m|\tilde{\beta}_P^r) = \tilde{\beta}_P^m \tilde{H}^*(\tilde{\beta}_P^r) - b - \gamma_P - \alpha_P(\tilde{\beta}_P^m) \tag{S14}$$

$$\tilde{\mathcal{F}}(\tilde{\beta}_P) = \frac{\partial \tilde{w}}{\partial \tilde{\beta}_P^m}(\tilde{\beta}_P^m|\tilde{\beta}_P^r)\bigg|_{\tilde{\beta}_P^m=\tilde{\beta}_P^r=\tilde{\beta}_P} = \tilde{H}^*(\tilde{\beta}_P) - \frac{\mathrm{d}\alpha_P}{\mathrm{d}\tilde{\beta}_P}(\tilde{\beta}_P). \tag{S15}$$

The value of $\tilde{\beta}_P$ that satisfies $\tilde{\mathcal{F}}(\tilde{\beta}_P) = 0$ is called our singular strategy, and is calculated as:

$$\tilde{\beta}_P^* = \left(\frac{b + \gamma_P + \bar{\alpha}_P}{(d-1)\bar{\alpha}_P}\right)^{1/d}. \tag{S16}$$

We can show that this is a continuously stable strategy (CSS) by demonstrating that it is both evolutionary stable (ES) and convergence stable (CS). Evolutionary stability requires the second derivative of $\tilde{w}$ with respect to the mutant strain being negative:

$$\frac{\partial^2 \tilde{w}}{\partial (\tilde{\beta}_P^m)^2}(\tilde{\beta}_P^m|\tilde{\beta}_P^r) = -\frac{\mathrm{d}^2\alpha_P}{\mathrm{d}(\tilde{\beta}_P^m)^2}(\tilde{\beta}_P^m). \tag{S17}$$

Since the virulence of the parasite is an accelerating function, the second derivative of the virulence function is positive and hence the singular strategy is evolutionary stable. For convergence stability, we need to differentiate the fitness function once with respect to the mutant and once with respect to the resident, and ensure that this is larger (S17):

$$\frac{\partial^2 \tilde{w}}{\partial \tilde{\beta}_P^m \partial \tilde{\beta}_P^r}(\tilde{\beta}_P^m|\tilde{\beta}_P^r) = \frac{\partial \tilde{H}}{\partial \tilde{\beta}_P^r}(\tilde{\beta}_P^r). \tag{S18}$$

Note that at the singular strategy, we have $\tilde{H}^*(\tilde{\beta}_P^*) = \alpha'_P(\tilde{\beta}_P^*)$ (from (S15) equalling 0). Then

$$\begin{aligned}\left.\frac{\partial^2 \tilde{w}}{\partial \tilde{\beta}_P^m \partial \tilde{\beta}_P^r}(\tilde{\beta}_P^m|\tilde{\beta}_P^r)\right|_{\tilde{\beta}_P^m=\tilde{\beta}_P^r=\tilde{\beta}_P^*} &= \left.\frac{\partial \tilde{H}}{\partial \tilde{\beta}_P^r}(\tilde{\beta}_P^r)\right|_{\tilde{\beta}_P^r=\tilde{\beta}_P^*} \\ &= \frac{\alpha'_P(\tilde{\beta}_P^*) - \tilde{H}^*(\tilde{\beta}_P^*)}{\tilde{\beta}_P^*} \\ &= 0.\end{aligned}$$

Therefore, the singular strategy is a CS and hence is a CSS.

#### 3 Definition of strong convergence stability

In order to determine if any co-singular strategies are convergence stable, we use the definition presented by Leimer (2009). Suppose we have fitness functions for the defensive symbiont and the parasite defined by $w_D(y^m|\boldsymbol{\theta}^r)$ and $w_P(\beta_P^m|\boldsymbol{\theta}^r)$ respectively, where $\boldsymbol{\theta}^r$ defines the resident traits, and trait values with a superscript $m$ denote the invading mutant trait. The expressions for each of these are given in the main text (equations (6) and (7)). Suppose that we have a singular strategy, which we write as $\boldsymbol{\theta}^* = (y^*, \beta_P^*)$. Then this singular strategy is *strong convergence stable* if the following three conditions hold:

$$\left(\frac{\partial^2 w_D}{\partial (y^m)^2} + \frac{\partial^2 w_D}{\partial y^m \partial y^r}\right)\left(\frac{\partial^2 w_D}{\partial (\beta_P^m)^2} + \frac{\partial^2 w_D}{\partial \beta_P^m \partial \beta_P^r}\right) > \frac{\partial^2 w_D}{\partial y^m \partial \beta_P^r}\frac{\partial^2 w_P}{\partial \beta_P^m \partial y^r}, \tag{S19}$$

$$\frac{\partial^2 w_D}{\partial (y^m)^2} + \frac{\partial^2 w_D}{\partial y^m \partial y^r} < 0, \tag{S20}$$

$$\frac{\partial^2 w_D}{\partial (\beta_P^m)^2} + \frac{\partial^2 w_D}{\partial \beta_P^m \partial \beta_P^r} < 0, \tag{S21}$$

with all derivatives evaluated at the singular strategy.

Olaf Leimer. Multidimensional convergence stability. *Evolutionary Ecology Research*, 2009.
